# A Plasmid System with Tunable Copy Number

**DOI:** 10.1101/2021.07.13.451660

**Authors:** Miles V. Rouches, Yasu Xu, Louis Cortes, Guillaume Lambert

**Affiliations:** Field of Biophysics, Cornell University, Ithaca, NY 14853, USA; School of Applied and Engineering Physics, Cornell University, Ithaca, NY 14853, USA

## Abstract

Plasmids are one of the most commonly used and time-tested molecular biology platforms for genetic engineering and recombinant gene expression in bacteria. Despite their ubiquity, little consideration is given to metabolic effects and fitness costs of plasmid copy numbers on engineered genetic systems. Here, we introduce two systems that allow for the finely-tuned control of plasmid copy number: a plasmid with an anhydrotetracycline-controlled copy number, and a massively parallel assay that is used to generate a continuous spectrum of ColE1-based copy number variants. Using these systems, we investigate the effects of plasmid copy number on cellular growth rates, gene expression, biosynthesis, and genetic circuit performance. We perform single-cell timelapse measurements to characterize plasmid loss, runaway plasmid replication, and quantify the impact of plasmid copy number on the variability of gene expression. Using our massively parallel assay, we find that each plasmid imposes a 0.063% linear metabolic burden on their hosts, hinting at a simple relationship between metabolic burdens and plasmid DNA synthesis. Our plasmid system with tunable copy number should allow for a precise control of gene expression and highlight the importance of tuning plasmid copy number as tool for the optimization of synthetic biological systems.

## INTRODUCTION

Plasmids are an invaluable tool in biotechnology and genetic engineering due to their small size and the ease with which they can be engineered for biotechnological applications. Recent studies have highlighted the potential benefits of fine-tuning plasmid copy number to the task at hand. For example, runaway replication has been used as a method for increasing cellular protein production [1, 2]. Recent works have also leveraged plasmid copy numbers to amplify a signal through a large increase in copy number [3, 4] and to increase the cooperativity and robustness of synthetic gene circuits [5]. Single cell measurements have also been used to accurately quantify the prevalence of plasmid loss within a population [6].

While recent studies have probed the effect of plasmid copy number on host cell growth [7, 8] and the role of plasmids on central metabolic processes [9], we still lack a quantitative description of the relationship between plasmid copy number and host metabolic burden. Indeed, recombinant gene expression results in significant metabolic burden on bacterial cells [10–12] which may interfere with normal cellular processes and even lead to cessation of growth. The control of plasmid copy number can also drastically alter the behavior of simple repression systems [13, 14] which may significantly impact engineered gene circuits and should be important considerations for plasmid-based biotechnology and bioengineering applications

In this work, we use two novel approaches to actively control the copy number of pUC19 and other ColE1-based plasmids to study the impact of copy number on cellular growth and metabolism. Specifically, we first introduce a strategy for finely tuning plasmid copy number in a way that is dependent on the concentration of an exogenous inducer molecule. Then, we develop a method which uses a massively parallelized assay for rapid screening of the dependence of a system on plasmid copy number.

We further characterize our systems at both the macroscopic and single cell level to uncover a relationship between gene copy number, protein production, and cellular growth rates. Our approach also leads to insights into the variation in plasmid copy numbers and the phenomena of plasmid loss and runaway replication at the singlecell level. We finally demonstrate that manipulations of plasmid copy numbers can be used to tune the behavior of synthetic gene circuits such as a simple CRISPRi genetic inverter, and control the expression of complex biosynthetic pathways to optimize the production of the pigment Violacein. Our work thus provides a straightforward method for generating copy number variants from a specific plasmid backbone, which we anticipate will expand the synthetic biology toolkit and constitute the basis for more nuanced attention to plasmid copy number in synthetic gene circuits, protein biosynthesis, and other biotechnological applications.

## RESULTS

### Model of ColE1 Plasmid Replication and Copy Number

The molecular mechanisms that regulate plasmid copy number control have been well characterized [16]. In the ColE1 origin of replication, used extensively in this work, two RNAs control replication (Fig. 1A): the priming RNA (RNA-p), which acts in *cis* as a primer near the origin of replication, and the inhibitory RNA (RNA-i) which is transcribed antisense to RNA-p and acts in *trans* to inhibit replication through binding to the priming RNA prior to hybridization near the origin (Fig. 1B) [17].

**Fig. 1.**
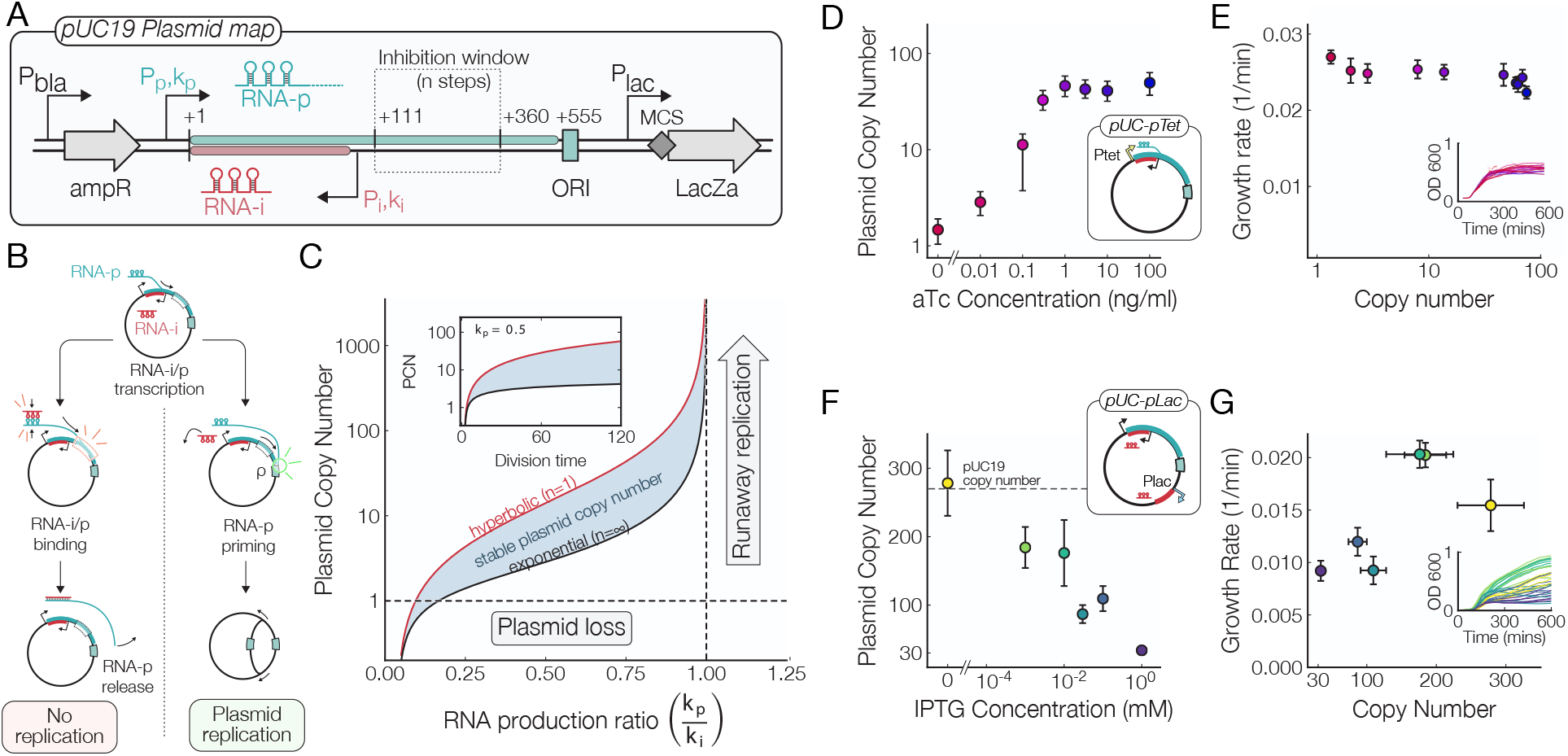
Replication of ColE1 Origin Plasmids. A) Map of the pUC19 plasmid. The origin of replication consists of a 555 bp region containing the priming RNA (RNA-p), the inhibitory RNA (RNA-i), the inhibition window, and the site of replication initiation (ORI). B) Diagram of the steps leading to ColE1 replication. In the ColE1 Origin, reversible binding of the inhibitory RNA-i to RNA-p exercises control over the copy number by inhibiting replication (left branch). Production of RNA-p transcript may result hybridization with the ORI, which leads to plasmid replication (right branch). C) Predicted plasmid copy number for hyperbolic (upper) and exponential (lower) initiation models [15] plotted as a function of RNA-p transcription initiation rate. Inset shows the predicted dependence of plasmid copy number on host cell division time. Model parameters: *ϵ* =0.545 / min, *ρ* = 0.65, *r* =0.01 / min, *k_i_* =0.95 / min. D) Plasmid copy number measurements made by digital droplet PCR of the aTc-inducible pUC-pTet plasmid at increasing aTc concentrations. E) Growth rates of cells hosting the pUC-pTet plasmid at increasing plasmid copy numbers. We see a marginal effect of plasmid copy number on cellular growth rates, inset shows individual growth curves colored by aTc concentration. F) Plasmid copy number measurements of a plasmid which contains an additional copy of the inhibitory RNA under the control of an IPTG-inducible promoter. G) Growth rate vs plasmid copy number of the IPTG-inducible plasmid shows a strong metabolic burden at high levels of RNA-i expression.

A simple macroscopic-scale model developed by Paulsson *et al*. recapitulates these observations and predicts the functional dependence of plasmid copy number on the molecular details of this process [15] (Fig. 1C). This process is classified as inhibitor-dilution copy number control [18], owing to the fact that the copy number is controlled through a combination of replication inhibition and dilution of plasmids and regulatory components due to cell division.

Specifically, as the ratio of RNA-p transcription rate *k_p_* to RNA-i transcription rate *k_i_* is increased, plasmid copy number *N_p_* will also increase according to

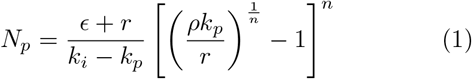

where *ϵ* is the RNA degradation rate, *ρ* is the RNA-p priming probability, *r* is the cellular growth rate, and *n* is the number of rate-limiting steps in the inhibition window [15]. The *n* → 1 and *n* → ∞ limits represent hyperbolic and exponential inhibition, respectively (Fig. 1C).

As the priming:inhibitory RNA production ratio *k_p_*/*k_i_* approaches 1, the number of inhibitory RNA will not be present in high enough number to sequester priming RNA transcripts, thereby failing to limit plasmids from duplicating in a process called runaway replication (Fig. 1C). In addition, increasing the plasmid copy number may increase the metabolic burden associated with plasmid expression and reduce the growth rate *r*, which in turn increases the plasmid copy number further as *r* → 0.

On the other hand, as the RNA production ratio *k_p_*/*k_i_* decreases, the number of plasmids inside each cell will approach 1 and small variations in the number of plasmids apportioned to each daughter cell during cell divisions may lead to the catastrophic consequence of plasmid loss. These two stresses, plasmid loss and metabolic burden, necessitate the evolution of copy number control systems to ensure that neither stress results in the loss of the plasmid [19].

### Tuning Priming and Inhibitory RNA Transcription to Control ColE1 Copy Number

The Paulsson model of ColE1-type replication predicts that the RNA-p transcription initiation rate should determine the plasmid copy number without drastically altering the sensitivity of copy number control [18]. Thus, in order to only control the activity of the promoter upstream of the priming RNA, we first replaced the native priming promoter in the pUC19 plasmid with an anhydrotetracycline (aTc) inducible promoter (Fig. 1D). Liquid cultures of *E. coli* overexpressing TetR and carrying this plasmid were grown in aTc concentrations spanning 4 orders of magnitude and the plasmid copy number was determined via digital droplet PCR (ddPCR) from total DNA isolates by measuring the ratio of the *bla* gene on the plasmid to the genomic *dxs* gene [20].

This plasmid showed robust induction of copy number with aTc induction (Fig. 1D). Copy numbers ranged from 1.4 copies per cell in the complete absence of aTc to roughly 50 copies per cell at full induction (100ng mL^−1^ aTc). We saw a very similar relation in the plasmid yield from overnight cultures grown at varying aTc concentrations (Sup.).

We subsequently measured the growth rate of cells harboring the pUC-pTet plasmid as a function of the aTc concentration in a microplate reader (Fig. 1E). Though the absolute difference in growth rates is small, a cell with a few copies of the plasmid grows roughly 10% faster than a cell with 50 plasmids, we can clearly see that increasing the plasmid copy number has a weak, though consistent, effect on the host cell growth rate.

To complement the work above, we sought to demonstrate that the transcription level of inhibitory RNA can be used to control ColE1 copy number as well. Seeking to do this with minimal perturbation to the native pUC19 origin of replication, we inserted a second copy of the inhibitory RNA downstream of the IPTG-inducible Lac promoter on the pUC19 plasmid. This gives us control over the inhibition of replication through the addition of exogenous *IPTG* (Fig. 1F).

When no *IPTG* was added to the cells we measured ~270 plasmids/genome, a number that agrees with what we have measured for the standard pUC19 plasmid, indicating that our system efficiently represses the additional inhibitory RNA transcript and does not affect the copy number in the absence of induction. Induction of the inhibitory RNA greatly reduced plasmid copy number to approximately 30 plasmids/genome at full RNA-i induction, demonstrating that tuning RNA-i production rate can be used to control the plasmid copy number.

It is interesting to note that we observed paradoxical effects on the growth rate from cells harboring this plasmid - at high plasmid copy numbers we see a faster growth rate and at lower copy numbers we see a markedly slower growth rate (Fig. 1G). These observations are markedly different from what we see in the pUC-pTet plasmids, where increased plasmid copy number has a relatively modest effect on the growth rate of host cells. Instead, the reduction in the maximum cell density reached by the IPTG-induced populations (Fig. 1G, inset) suggests that overproduction of the inhibitory RNA may completely abolish plasmid replication in some cells, preventing them from dividing further.

### Gene Expression Scales with Plasmid Copy Number

Understanding the interplay between plasmid copy number, host cell growth rate, and gene expression is key to optimizing protein production in bacterial cell cultures. If too much protein is produced, cells may stop growing or be out competed by cells harboring plasmids that lack the correct insert. If protein production is too low, on the other hand, one may have to process massive amounts of cells to obtain a usable amount of their target protein. Understanding the interplay between plasmid regulation and protein expression is key to solve protein production maximization problems.

To directly measure the effects of plasmid and gene copy number on gene expression and host cell growth rate, we first inserted sfGFP downstream of a constitutive promoter on our pUC-pTet plasmid, whose copy number is controlled through exogenous aTc induction. This gives us a straightforward way of monitoring the effects of plasmid copy number on gene expression. Cell cultures with this tunable copy number plasmid were grown in a microplate reader where simultaneous measurements of fluorescence and growth rate could be made.

We first observe clear differences in growth rates as a function of both plasmid copy number and aTc concentration (Fig. 2B). We see a roughly 15% decrease in host growth rate at maximal induction of copy number (~ 50 copies per cell). This metabolic burden is a significantly higher than the pUC-pTet plasmid (Fig. 1E), indicating that the majority of the reduction in growth rate comes from the overexpression of GFP and not from the increased plasmid copy number. In this sense, addition of GFP to the plasmid provides a stronger coupling between the host growth rate and plasmid copy number by taking up additional cellular resources.

**Fig. 2.**
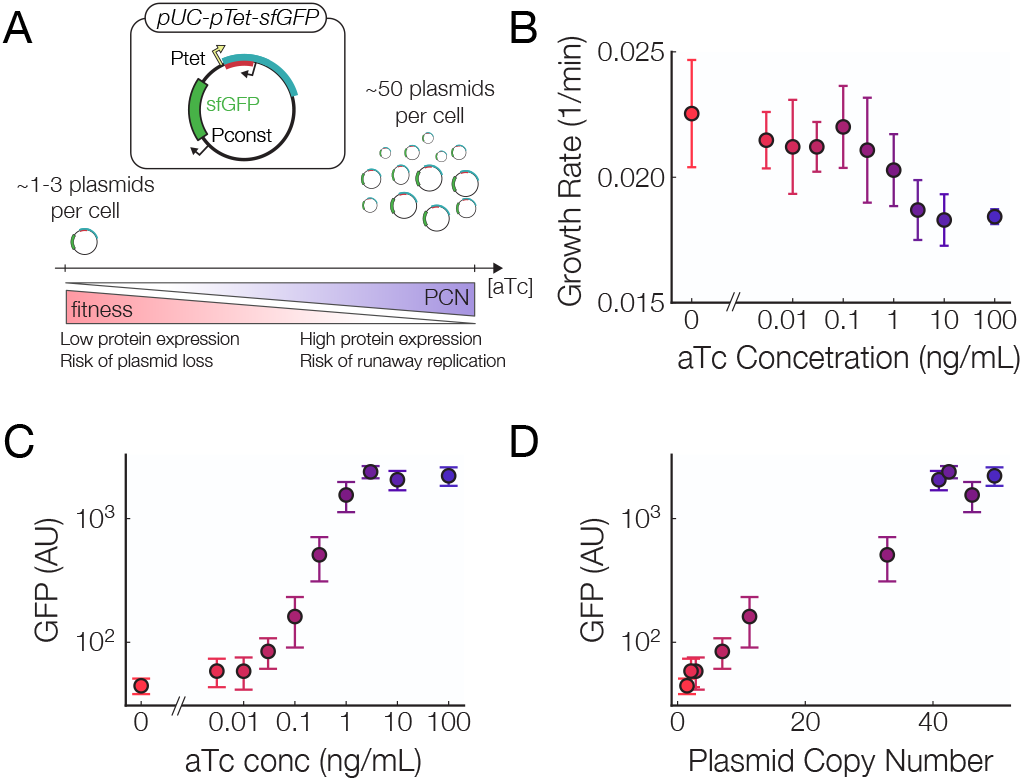
Control of Gene Expression through Induction of Plasmid Copy Number. A) Schematic of the effects of adding aTc to the pUC-pTet-GFP plasmid. As the concentration is increased the number of plasmids and GFP genes increases as well as the metabolic cost of the plasmid. B) Growth rates of cells hosting the pUC-pTet-sfGFP plasmid as a function of aTc concentration. C) GFP Production of the pUC-pTet-GFP plasmid grown at different aTc concentrations. D) GFP production of the pUC-pTet-GFP plasmid as a function of plasmid copy number as measured by digital droplet PCR.

Despite the fitness costs associated with increasing the plasmid copy number, we nevertheless observe an almost perfect GFP induction curve when compared to the aTc concentration (Fig. 2C). In addition, Fig. 2D shows a nearly linear increase in the fluorescence with the plasmid copy number, suggesting that the metabolic costs associate with protein overproduction only marginally impact the overall GFP production rate. Our pUC-pTet plasmid provides a simple way to control gene expression without the need to insert an inducible promoter upstream of the gene of interest.

### Metabolic Costs of the pUC-pTet Plasmid at the Single-Cell Level

While macroscopic measurements show a clear interdependence between gene expression and copy number, single cell measurements can provide a more powerful path towards understanding the relationship between plasmid copy number and gene expression. In particular, singlecell studies allow us to determine the distributions of gene expression of single cells at varying plasmid copy numbers and directly observe the phenomena of plasmid loss and runaway replication.

To first corroborate protein production measurements with plasmid copy number, we observed the pUC-pTet-sfGFP system for up to 150 generations in a microfuidic device [21, 22].

In agreement with population-wide measurements (Fig. 2C), individual cells show an increase in the mean fluorescence with plasmid copy number (Fig. 3A). We also observe a marked increase in the fluorescence of some cells at high aTc concentrations. Interestingly, cells with an increased fluorescence level are found to be elongated (Fig. 3A and Fig. S2). The intensity distribution (Fig. 3B) undergoes a profound change as the plasmid copy number increases: the distribution is peaked at a very low intensity at low plasmid copy number but, as the copy number increases, the distribution widens and a high-intensity tail emerges.

**Fig. 3.**
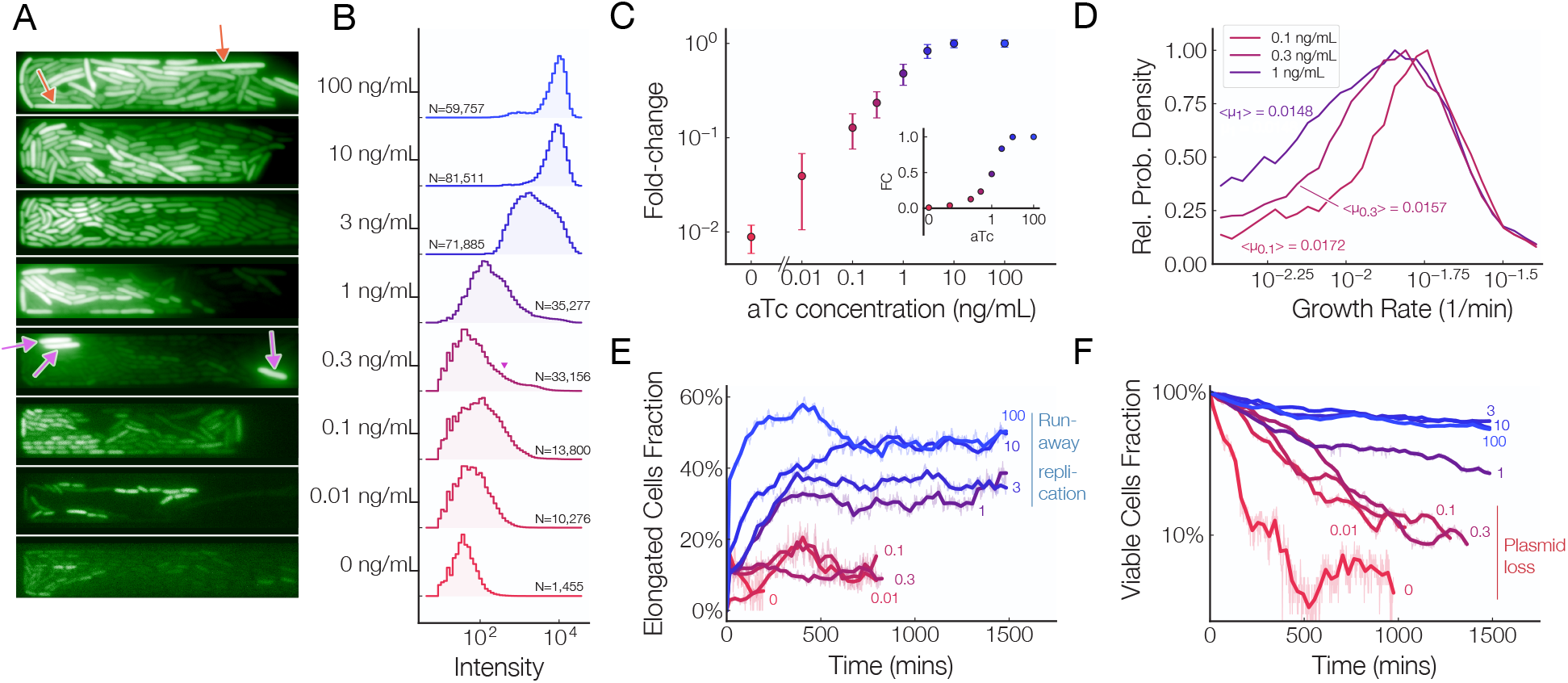
Microscopic Analysis of sfGFP Expressed by the pUC-pTet. A) Representative snapshots of microfluidic chambers at various aTc concentrations. Slow-growing cells at high copy number are indicated by orange arrows and cells at 0.3ng mL^−1^ of aTc in an “activated” state are indicated by purple arrows. B) Distribution of cellular fluorescence intensities at increasing aTc concentrations. C) Normalized mean fluorescence as a function aTc concentration, log scale. Inset: Normalized mean fluorescence vs aTc concentration on a linear scale. D) Growth rate distributions of cells growth at 0.1, 0.3 and 1ng mL^−1^ aTc. E) Fraction of slow-growing, elongated cells over time at several aTc concentration. This measurement is indicative of the extent to which cells are burdened by the high plasmid copy numbers. F) Proportion of viable cells as a function of time at various aTc concentrations. The sharp decrease in the number of viable cells at low aTc concentrations provides insight into the extent to which plasmid loss occurs at low copy numbers.

The most drastic shift in the distribution occurs between 0.3 ng mL^−1^ and 1 ng mL^−1^, where the *average* cell intensity (Fig. 3C) is in good agreement with population wide measurements (Fig. 2C), but each population sees a steady increase in the number of *individual* cells with increased GFP levels (Fig. 3A, red arrows). As the plasmid copy number increases further, a growing number of cells reach an “activated” state where they show fluorescence levels as high as those full pTet induction. Eventually, all cells for aTc concentrations above 3ng mL^−1^ reach this activated, fully-induced state (Fig. 3C).

As the plasmid copy number sharply increases between 0.1 ng mL^−1^ and 1ng mL^−1^ of aTc, we observe a clear drop in average growth rate (Fig. 3D and Fig. S4) similar to bulk measurements (Fig. 2B). The distribution of cellular growth rates (Fig. 3D) show that this drop is largely due to a fraction of cells that suffer from reduced growth rate. This is likely due to the metabolic burden of plasmid maintenance and GFP expression severely limiting the growth of a subset of the cellular population. Above 1ng mL^−1^ aTc, the growth rate of healthy cells seems to stabilize (Fig. S4). However, we notice an increasing fraction of elongated cells (Fig. 3E), many of which have lost ability to grow and turned extremely bright (supplementary movie M1). The inability of a fraction of cells to grow may explain why growth rate continually decreases even though plasmid copy number saturates at high aTc levels in bulk measurements (Fig. 2B).

At low plasmid copy numbers, we see a decreased number of cell in each microfluidic chamber. In fact, nearly every cells undergoes lysis within the first 1500 min of tracking for aTc concentrations lower than 1ng mL^−1^, suggesting that cells may not be resistant to carbenicillin due to plasmid loss (Fig. 3F).

These observations highlight the delicate balance between plasmid loss at low copy number and heightened metabolic burdens at high copy numbers. Our results demonstrate that both phenomena occur stochastically and on a per-cell basis.

### Massively Parallelized Assay Determines Copy Numbers of ColE1-derived Plasmids

Though we were able to obtain robust control over plasmid copy number with our inducible plasmids, we next sought to exert control over plasmid copy number without the need for external inducer molecules. In order to survey the landscape of possible plasmid copy numbers that arise from changes of the transcription rates of the RNAs controlling ColE1 replication, we simultaneously constructed 1024 different variants of the pUC19 plasmid through site directed mutagenesis of the −35 and −10 boxes of the promoter controlling the priming RNA and at the +1 site of the priming RNA transcript (Fig. 4A, Pp library). The same procedure was performed on the promoter controlling the inhibitory RNA (Fig. 1A, Pi library), giving us 2 separate libraries of plasmids: one with a variable priming RNA transcription rate *k_p_* and another with a variable inhibitory RNA transcription rate *k_i_*.

**Fig. 4.**
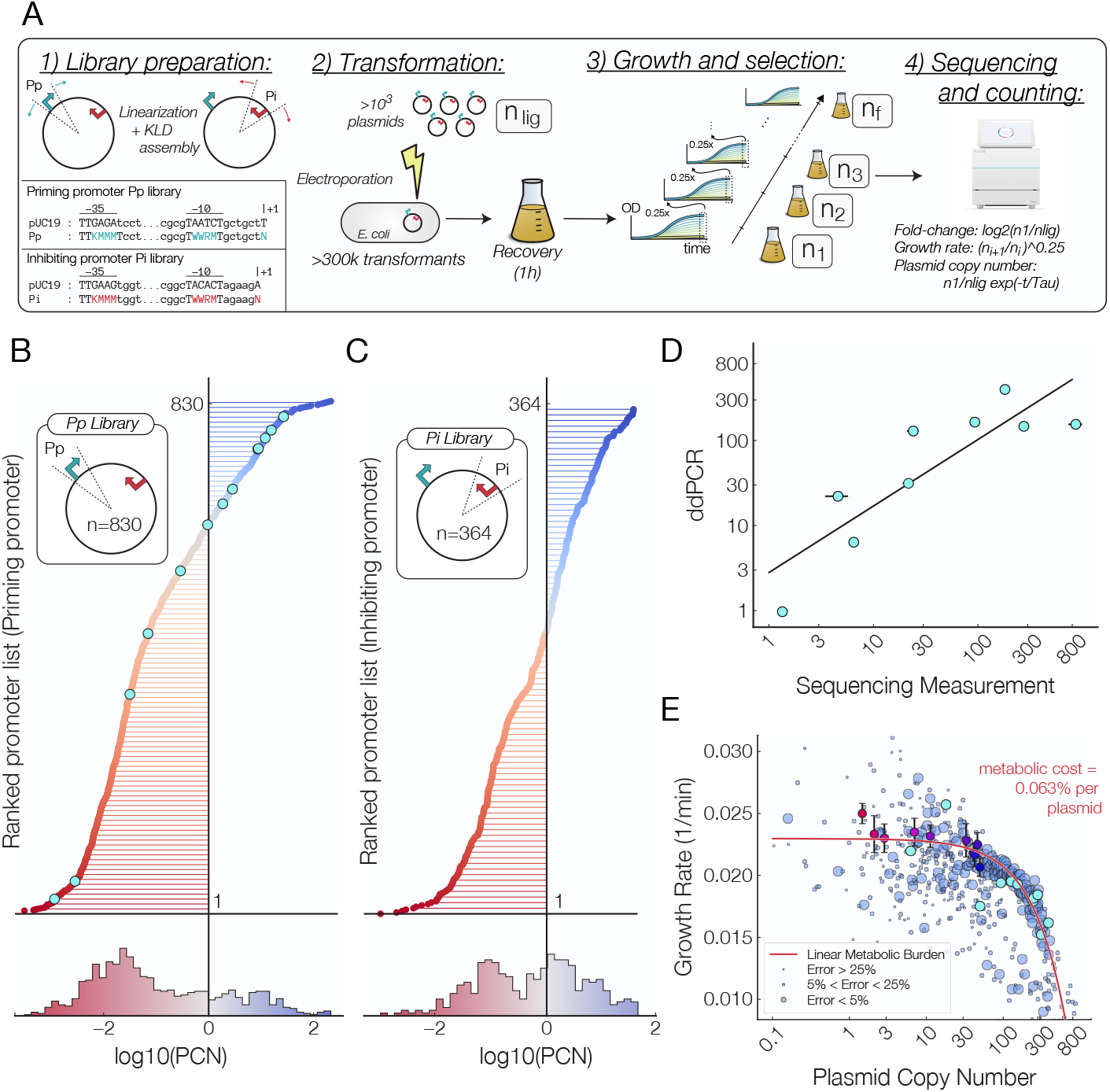
Massively Parallelized Copy Number Assay. A) Procedure for massively parallelized copy number assay. The promoter upstream of the priming RNA was randomized with site directed mutagenesis. 2. Plasmids were ligated and cells were transformed with this library of plasmids with high efficiency and grown until OD_600_ of 1.0, 3. at which point a fraction of the culture was diluted and regrown and the remaining fraction was saved for plasmid extraction. 4. Sequencing libraries were generated from isolated plasmids and sequenced using next generation sequencing. Growth rates and Copy numbers were then calculated from the frequencies of sequencing counts present at each time point. Digital PCR was performed to measure the copy number of individual constructs so that absolute copy numbers could be estimated from sequencing measurements. B) Ranked list of plasmids generated through mutations to the promoter driving the priming RNA in this work (*n* = 830). (lower) The distribution of copy numbers arising from mutations to the priming RNA promoter. C) Ranked list of plasmids generated through mutations to the promoter driving the inhibitory RNA in this work (*n* = 364). D) Agreement between plasmid copy number as measured by sequencing counts and digital droplet PCR. E) The relationship between plasmid copy number and growth rate as measured across 830 variants of the pUC19 plasmid. The size of each data point is inversely proportional to the error in the measurement, with the top, middle two and lowest quartiles representing an error of less than 5%, between 5% and 25%, and more than 25%, respectively. Data from Fig. 1E is plotted (red to blue gradient points) A linear metabolic burden seems to account for the observed relation reduction in growth rate with increasing copy number.

These libraries were constructed using multiplexed PCR and KLD assembly [23] and electroporated into competent *E. Coli* cells (Fig. 4A, step 2). Cell cultures were grown in selection conditions until they reached an OD_600_ of 1.0 and subsequently diluted by a factor of 4 and grown again to an OD_600_ of 1.0 (Fig. 4A, step 3). Excess culture from each step of dilution was set aside for sequencing (Fig. 4A, step 4).

For each recovered mutant, the abundance of each construct at each time point was used to infer the growth rate of each individual construct. By coupling this information with the fold change in sequencing counts from the library prior to transformation, we determined the relative copy numbers of every plasmid in this library (Figs. 4B-C) and their growth rates (Fig. 4D). Our method was able to generate 1,194 plasmids, each with a distinct plasmid copy number ranging from less than 1 copy per cell to close to 800 copies per cell (Table S1).

To make sure our sequencing-based method yields accurate plasmid copy number measurements, we next sought out to calibrate our sequencing-based numbers to absolute copy numbers made using digital droplet PCR measurements. Individual constructs of 9 promoters were selected (light blue datapoints in Fig. 4B) and their absolute copy number were determined via digital PCR by the ratio of *dxs* to *bla* genes [20]. There is a good agreement between fold change in sequencing abundance and the plasmid copy numbers as measured by digital droplet PCR (Fig. 4D), indicating that the fold change in abundance of sequencing counts with a modest correction applied to correct for differences in growth rates is a sound measure of plasmid copy number.

The simultaneous measurement of plasmid copy numbers and the growth rates of their host cells provides us with a powerful measurement of the interdependence of copy number and metabolic burden. While a high plasmid copy number should reduce host cell growth, to what extent that occurs is not known. Our measurements of plasmid copy numbers and growth rates for the 830 plasmids in the priming promoter library, all with nearly identical origins of replications, provide a powerful way to deduce this relationship.

In Fig. 4E, we observe that, as the plasmid copy number increases, the growth rate of host cells decreases. A simple relationship in which the metabolic burden of plasmid expression is linear with the copy number largely agrees with these findings –ie. cost = *a* · *b* · *x* + *b*, where *x* is the plasmid copy number, with *a* =0.063% and *b* =0.023 min^−1^. This relationship can arise from a simple resource allocation model in which each plasmid takes up a small fraction of the resources necessary for cellular growth proportional to the plasmid size (pUC19 is 2686 bp in size, which is approximately equal to 0.058% the length of the *E. coli* genome). This results opens up the possibility for improved models of plasmid evolution and replication, as they indicate that plasmids impose a metabolic burden that is linear with their copy number on their hosts. Knowledge of this relationship between host growth and plasmid copy number may shed light on the evolution of *cis* and *trans* acting regulatory elements as well as the stringency of plasmid copy number control mechanisms.

### Plasmid Copy Number Controls Violacein Biosynthesis

We sought to demonstrate the potential for using plasmid copy number to control complex biosynthetic processes comprising of several proteins and enzymes. We chose to focus on Violacein (Fig. 5A), a purple pigment formed by the condensation of two tryptophan molecules [24] which is commonly used to demonstrate biosynthesis optimization [25]. The violacein biosynthetic pathway contains five enzymes, VioABCDE, each under the control of a different promoter and spanning approximately 8kb of DNA (Fig. 5B). Hence, we reasoned that tuning the plasmid copy number of the whole Violacein production pathway may serve as an effective method to scale up the expression of VioABCDE while keeping the production of each enzyme under the same stoichiometric ratio.

**Fig. 5.**
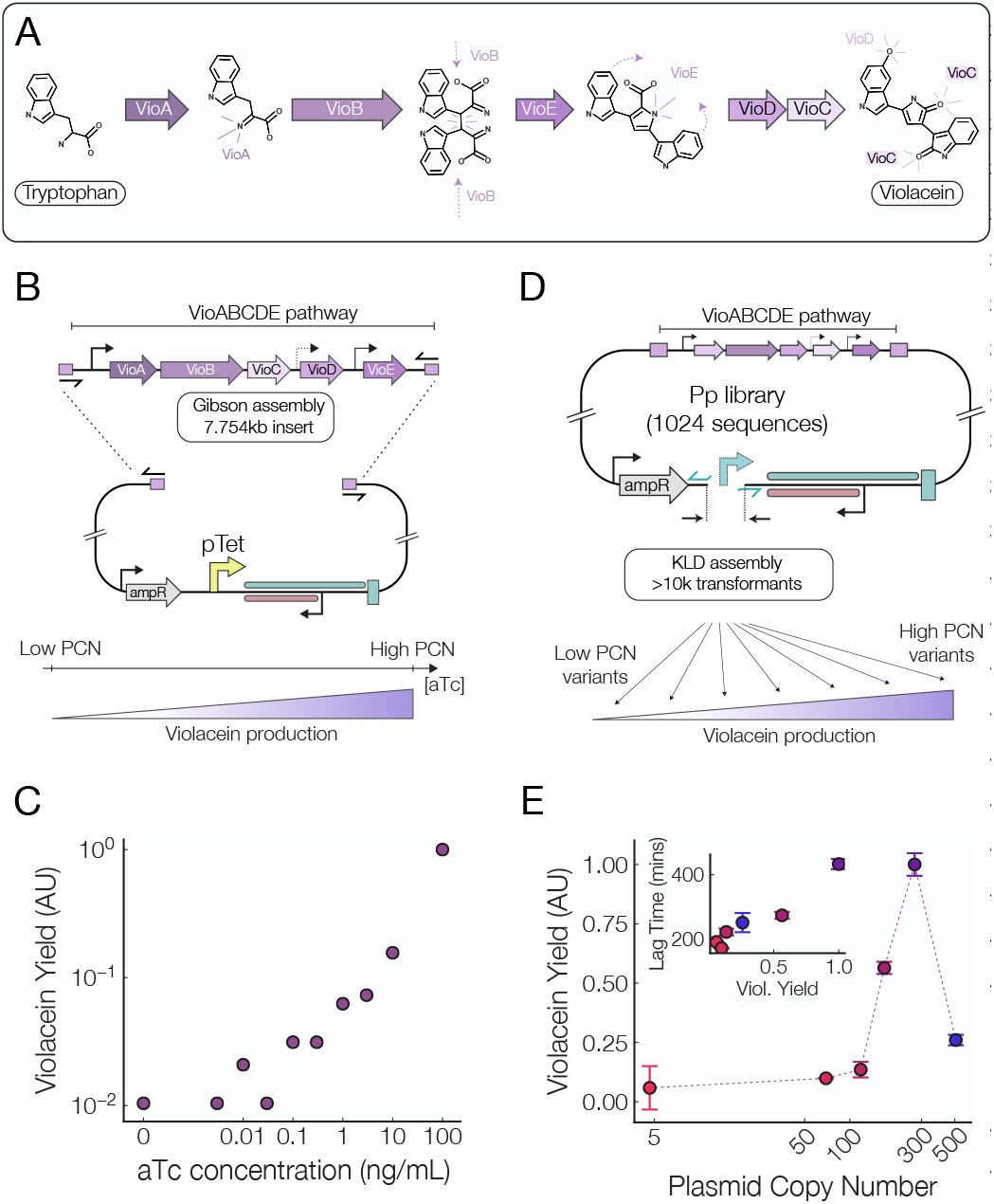
Optimization of Violacein Production by Tuning Plasmid Copy Number. B) The complete VioABCDE pathway was inserted in the pUC-pTet plasmid. C) Controlling violacein biosynthesis by increasing aTc concentrations. D) A library of variable copy number plasmids was created by performing mutagenesis on the priming promoter of a ColE1based plasmid containing the VioABCDE pathway. E) Violacein production peaks as the plasmid copy number nears 300, with a marked decrease in the yield as plasmid copy number is increased beyond 500. Inset: relationship between the violacein yield and lag time of the population when grown in a plate reader.

To achieve this, constructed a pTet inducible plasmid harboring the Violacein biosynthesis pathway using Gibson assembly [26] (Fig. 5B). We then grew cells with the pUC-pTet-Vio plasmid at increasing aTc concentrations and observed a clear increase in the violacein production with increasing aTc concentrations (Fig. 5C).

Additionally, we inserted the VioABCDE pathway into a pUC plasmid under the control of a moderate-strength promoter, and we used site directed mutagenesis of the origin of replication, in the manner described above, to generate a plasmid copy number library (Fig. 5D). After this library was transformed into competent cells, we picked individual colonies from a plate and sequenced their plasmids in order to infer their copy number from the Pp library data (Table S1). We also measured the violacein production from cell extracts and found a clearly positive relation between the plasmid copy number and Violacein production (Fig. 5E). Interestingly, we observed a marked decrease in violacein yields as the plasmid copy number is increased above 500 copies per cell, and growth curve measurements performed using a plate reader show that the decreased yield can be attributed to a longer lag time (Fig. 5E, inset). Our results show that using tunable copy number thus constitutes a generalizable method for the scaling-up and optimization of complex biosynthetic pathways.

### Control of a CRISPRi system through sponge binding sites

Controlling the plasmid copy number may also be useful in synthetic biology and in the creation of robust gene circuits [3, 5, 27, 28]. To test this, we used a simple CRISPR-dCas12a inverter to demonstrate the effect of adding “sponge” binding sites to a simple genetic circuit through manipulation of plasmid copy numbers. Sponge sites are additional binding sites used to titrate out (“soak up”) transcription factors, and they have been used to redirect flux in metabolic pathways to optimize arginine production [29], to activate silent biosynthetic gene clusters [30], to tune gene expression timing [31], and mitigate protein toxicity [32]. In these studies, variable numbers of sponge sites are placed on high or low copy number plasmids.

Here, we added a sponge site to a few isolates from our tunable copy number library to alter the qualitative behavior of a simple genetic circuit. Specifically, we use a two plasmid system in which a low-copy plasmid (pSC101, 2-5 copies per cell) contains an aTc inducible CRISPR-Cas12a guide RNA which can bind to an active site *S_a_* and turn OFF a promoter driving sfGFP, while the second plasmid contains a decoy binding site *S_d_* site that sequesters CRISPR-Cas12a molecules away from the active site *S_a_*. The sponge plasmids were picked out from our library of variable copy number plasmids generated in Fig. 4B and span copy numbers from 30 to 270.

Time-series measurements of the fully-induced CRISPRi inverter show that the OFF state of the inverter has a higher fluorescence level as the number of sponge sites in increased (Fig. 6A). This suggests that the addition of sponge sites reduces the occupancy of the active site *S_a_* and decreases the repression efficiency of the CRISPRi inverter.

**Fig. 6.**
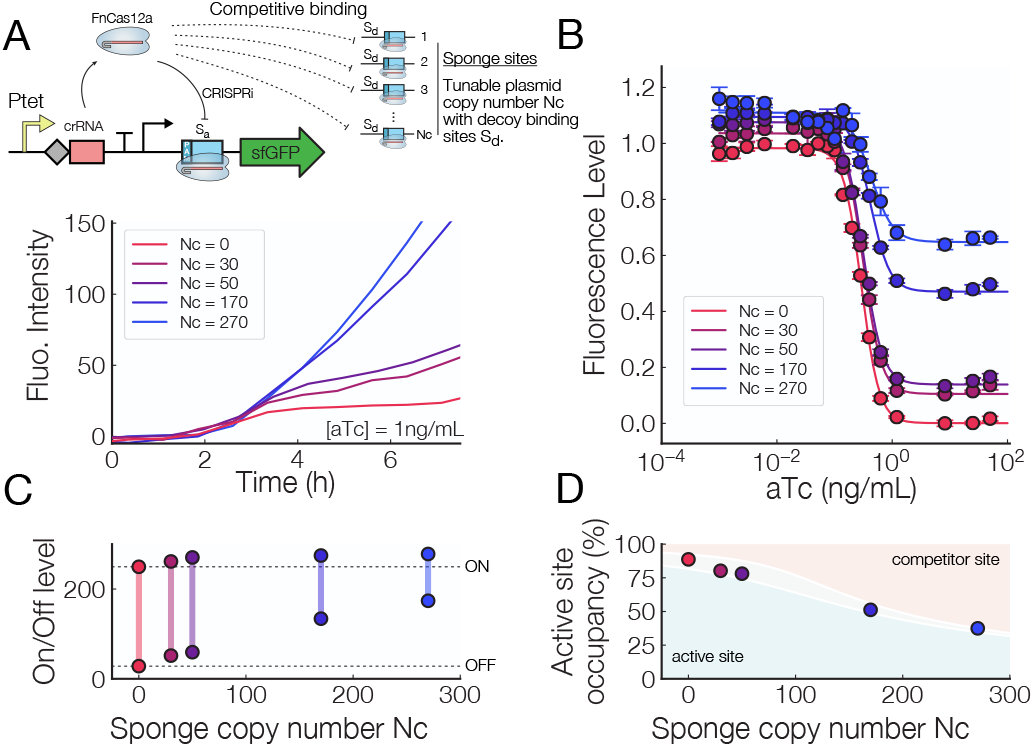
Control of a CRISPRi Inverter with Competitor Binding Sites. A) Top: Diagram showing how production of a dCas12a CRISPR RNA (crRNA) is controlled by an aTc-inducible promoter, and the dCas12a+crRNA duplex can either bind to the active site *S_a_*, which turns off production of sfGFP, or to a decoy site *S_d_* residing on a “sponge” plasmid. Bottom: Fluorescence intensity vs time for CRISPR-dCas12a systems with variable numbers of sponge sites (Nc). B) Induction curves for a CRISPRi system with sponge sites on plasmid with varying copy numbers. C) On-Off levels for variable numbers of sponge sites. Both the ON and OFF fluorescence level of the inverter increase as the number of sponge sites increases. The dynamic range decreases with increasing sponge copy number. D) Active site occupancy vs number of sponge sites, as the number of sponge sites increases the active site becomes less occupied. The gray area represents solution to the fold-change model [14] for a dCas12a binding energy equal to −13.25 k_B_T (top) and −12.25 k_B_T (bottom).

We also observe a drastic reduction in the dynamic range of circuits as the number of sponge sites is increased (Fig. 6B), where cells with more than 150 sponge sites have at least a 40% reduction in dynamic range compared to those with no sponge sites. In contrast, the addition of 30 to 50 sponge sites only reduces the dynamic range by approximately 6-8% (Fig. 6C).

While a large dynamic range is often desirable in genetic devices, a reduction in dynamic range could be used to divert the flux in a metabolic pathway, ensure that the concentration of some protein stays within a given range, or to reduce the toxicity of CRISPR-Cas proteins [33] due to spurious off-target binding. Using a simple model of protein-DNA binding [14], we can determine that the occupancy of the active binding site *S_a_* (Fig. 6D) in the OFF state is reduced from nearly 100 % with zero sponge sites to 50 % when 270 sponge sites are present. Overall, we demonstrate that plasmid copy number can be used to accurately tune the level of transcription factor titration in a simple genetic inverter.

## DISCUSSION

This work details the development of a system of tunable copy number plasmids and demonstrates their utility in protein expression, biosynthetic pathway optimization, and synthetic biology applications. Using a combination of high-throughput measurements and single-cell experiments, we deduce relations between plasmid copy number and host cells growth rate, protein expression, biosynthesis, and the performance of simple regulatory elements. This system could help resolve known issues currently plaguing the design and scaling up of synthetic gene circuits such as leaky expression, narrow ranges of activation and repression control, and a large metabolic burden due to competition over a common pool of resources [34–36].

Using a massively parallel directed mutagenesis approach, we find that the most straightforward and reliable way to change the plasmid copy number in the ColE1 Origin is through mutations to the promoter controlling the priming RNA. Further, changes to the machinery controlling inhibitory RNA levels are much less likely to give stable constructs given the fact that mutations to the promoter controlling inhibition change the sequence of the priming transcript can have drastic effects on plasmid copy numbers through secondary structure changes [37]. Overall, these results suggest that the ColE1 replication machinery is more robust to changes in the priming RNA transcription initiation rate and much less robust to changes in the parameters detailing inhibition.

Our multiplexed method of generating a plasmid copy number library allows one to rapidly screen the effects of plasmid copy number on a given system and could be tailored to select for genetic devices that perform the best under a given set of conditions. A key finding of this facet of the work is that ColE1-type plasmids impose a linear metabolic burden on their hosts proportional to the relative size of the plasmid, which may shed light on the evolution of various regulatory elements in inhibitor-dilution copy number control mechanisms.

Our single cell measurements also show that plasmid loss and runaway replication are quite common phenomena. These findings are in line with recent single cell measurements of plasmid copy numbers [6] and illustrate that very careful attention should be given to the plasmids used to house genetic circuits in order to ensure reliability. Since we observed an increasingly large fraction of non-growing cells as we increased the plasmid copy number, our results further suggest that lower growth rates population carrying a high-copy number plasmid may be explained by non-growing cells as opposed to a universal reduction in cellular growth rates. While this work is restricted to the ColE1 origin of replication, similar principles can be used to tune the copy numbers of other inhibitor-dilution controlled plasmids.

The implications of plasmid copy number go beyond its effects on gene expression and host cell fitness that we’ve presented thus far. As vehicles of horizontal gene transfer, the prevalence of which increases with plasmid copy number, plasmids play a key role in bacterial ecology and evolution[38]. On the other hand, a consequence of a high copy number is a reduced retention and fixation of novel plasmid variants [39]. Together, these two issues create a complicated image of the role of plasmid copy number in bacterial diversity and evolution, it’s possible that the methods for controlling plasmid copy number presented here can be used help shed light on the role of plasmid replication in bacterial evolution.

The evolution of the fitness cost associated with plasmid-borne antimicrobial resistance genes is of central importance in preventing the spread of drug-resistant bacteria [40]. Our work presented here could provide a clear path for investigating the role of plasmid copy number in the evolution of the cost of antimicrobial resistance. Further, the fitness cost of plasmid expression is highly species-dependent and dependent on the diversity of a bacterial community [38] and recent findings have shown that plasmids can quickly achieve fixation in a population under non-selective conditions [41], further understanding of plasmid replication mechanisms and copy number effects might allow for a more nuanced understanding of these varied properties of plasmids.

Together, these findings demonstrate that plasmid copy number can have drastic effects on host cellular processes and provides straightforward ways of investigating these effects in other systems. Our results underscore the importance of tuning plasmid copy number as tool for the optimization of biosynthetic pathways, gene regulatory circuits, and other synthetic biological systems.

## MATERIALS AND METHODS

### Next-generation sequencing library preparation

The multiplexed libraries of plasmids was generated by site-directed mutagenesis using the NEB Q5 High-fidelity polymerase. Primers (pBR322-promoter1-1024-rev 5’-GAKKKMAAGA AGaTcCTTTG aTcTTTTCTA CGGGGTCTG-3’,pBR322-promoter1-1024-for 5’-CTTTTTTTCT GCGCGTWWRM TGCTGCTNGC AAACAAAAAA ACCACCG-3, pBR322-promoter1-1024-rev 5’-GTTAGGCCAC CAKKKMAAGA ACTCT-GTAGC ACCGCCTACA TACC-3’, pBR322-promoter1-1024-for 5’-TACGGCTWWR MTAGAAGNAC AG-TATTTGGT aTcTGCGCTC TGCTG-3’) containing degenerate nucleotides in the −35, −10, and +1 sites of the promoters controlling transcription of both RNAs were used to amplify the standard pUC19 plasmid (NEB). After PCR amplification samples were digested with DpnI (NEB) restriction enzyme to remove excess pUC19 template.

The PCR product was then circularized with electroligase (NEB) and transformed with high efficiency (≥ 1.5 million CFU/mL) into Endura Electrocompetent cells. The transformation efficiency was determined by plating of serial dilutions of recovered cells. After transformation liquid cultures were grown in Terrific Broth (TB, VWR) until reaching an OD_600_ of 1.0, at which point cultures were diluted 1:4 and regrown to OD_600_ of 1.0, a procedure that was repeated 4 times. Excess culture from each time point was set aside for plasmid isolation. Plasmids were isolated by miniprep (Zymo) and a 243 bp region of the ori was amplified (pBR322-5prime-promoter1-rev 5’-TCTACACTCT TTCCCTA-CAC GACGCTCTTC CGaTcTCAGA CCCCGTA-GAA AAGaTcAAAG GaTcTTC-3’, pBR322-3prime-for 5’-GACTGGAGTT CAGACGTGTG CTCTTCCGAT CTCGAGGTAT GTAGGCGGTG CTACA-3’) via PCR, at which point universal primers and indices were attached to the segment using an additional PCR step (primers). Prepared libraries were then diluted to 50 pM and sequenced on the Illumina iSeq using overlapping 2 × 150 bp overlapping reads.

### Construction of inducible plasmids

The pUC-pTet plasmid, whose copy number is inducible in the presence of aTc was constructed using a PCR-based insertion as follows. Primers (pTet-tn10-for 5’-AAATAACTCT aTcAATGATA GAGTGTCAAG AAGaTcCTTT GaTcTTTTCT ACGGGGTCTG A3’, pTet-tn10-rev 5’-TACCACTCCC TaTcAGTGAT AGAGATaTcT GCAAACAAAA AAACCACCGC TACCAGC-3’) overlapping with the pUC19 origin of replication and containing a pTet-inducible promoter were used to linearize and amplify the pUC19 backbone. The linearized plasmid was then circularized using a KLD reaction (NEB) and transformed into competent cells via electroporation. During the recovery period 1x aTc was added to the recovery media to facilitate replication of the plasmid.

The pUC-pLac-RNAi plasmid, whose copy number can be reduced through the addition of IPTG, was constructed as follows. The inhbitor RNA (commonly called RNAI) from the pUC19 plasmid was isolated via PCR (primers), in the process 15 bp overhangs were added to facilitate Gibson assembly. Another pUC19 plasmid was the linearized downstream of the pLac promoter (primers) and the linearized plasmid together with the inhibitory RNA insert were assembled via Gibson assembly at 50°C for 1 h and transformed into competent cells via electroporation.

### Copy number and growth rate measurement via next-generation sequencing reads

To determine the relative plasmid copy number from the abundance of next generation sequencing reads we compare the fraction of the sequencing reads at several time points with that in the initial library. The ratio of the two, with a small correction applied to account for differences in the growth rates of cells harboring different plasmids, is what we report as a relative plasmid copy number.

We measure initial fraction of the reads in the ligation product (fi) and the final fractions of the reads of the plasmids harvested at different time points (ff,f2,f3,f4). Differences in the doubling time () can be measured from differences in the fractions of reads from points separated in time. These relationships can be described in the equations below. First, to determine the doubling time from adjacent time points:

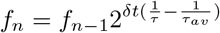

The above line states that the fraction at a given time point is equal to the fraction at the previous time point multiplied by the difference in the number of doublings from the bulk culture. Below we have rearranged the equation to solve for the growth rate from the variables we can either observe or control.

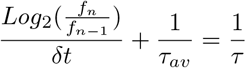

Secondly, to determine the copy number from the growth rate and the difference in sequencing counts we express the fraction at the first time point as the product of the fraction in the ligation, the relative copy number, and the number of excess doubling times:

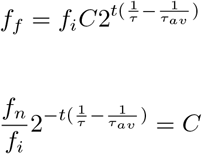

By inserting the above expression for the doubling time, we can have the relative copy number solely in terms of the fractions of each sequencing reads and the ratio of the time between time points and the time between transformation and the first time point.

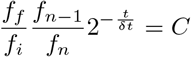

This analysis hinges on the assumption that upon transformation each cell receives a single plasmid, that cells do not transfer plasmids between each other, and that the number of transformants of a given plasmid is proportional to the amount of that construct present in the initial electroligation product. Further, this analysis assumes that cells harboring different constructs have identical lag times.

### Absolute Copy Number Determination via digital droplet PCR (ddPCR)

To make absolute determinations of plasmid copy numbers we used digital droplet PCR. Colonies hosting the appropriate constructs were picked off of plates after transformation and grown in 3 mL of TB or H media (10gL^−1^ tryptone, 8gL^−1^ NaCl) to OD_600_ of 1.0. Cells were then serially diluted 8000-fold in nuclease free ddH2O. Cells were lysed by heating at 95°C for 20 min. After lysis 1 μL of the lysate was used in each digital PCR reaction. Two digital PCR reaction were performed per sample - one targeting the genomic *dxs* gene and another targeting the beta lactamase, *bla*, gene on the plasmid (primers). This defines the plasmid copy number as the number of plasmids per genome. After generating droplets on the QX200 droplet generator (Biorad), reactions were thermal cycled at 95°C for 10 min, followed by 35 cycles of 98°C for 30 s, 58°C for 30 s, and 72°C for 1 min, the a 1 min extension at 72°C. Samples were held at 4°C after the reaction was completed and before droplets were read. Droplets were read with the QX-200 Droplet Reader (Biorad) and concentrations were determined using the Quantasoft analysis pro software (Biorad).

### PCN Sponge Effects on CRISPRi Inverter

CRISPR system with nuclease activity deactivated Cas protein(dCas) can be used to interfering transcription regulation(CRISPRi) by design crRNA targeting at the promoter for the gene of interest which acts a repressor of gene. However, dCas/crRNA complex can also bind to other site that shares the same DNA sequence as the target but not directly regulating the gene(competitor site). When the second scenario happens, it will have an impact on the repression level of the gene at given concentration of dCas/crRNA, i.e., a sponge effect for the gene of interest by consuming dCas/crRNA from the same pool. To test the sponge effect at different number of competitor sites, a dual-plasmid system was used. The main plasmid(Kanamycin resistant) is a low copy(pSC101) pTet-inducible CRISPRi-dFnCas12a inverter, where the crRNA transcribes by pTet target at a promoter that drivers a green fluorescence protein(sfGFP). The second plasmid is edited from the above mentioned pUC19-PCN library by inserting a single competitor via standard Site-Directed Mutagenesis(NEB, E0554). Two plasmids were co-transformed into GL002 strain that has dFnCas12a and TetR integrated in its genome with Mix & Go! E.coli Transformation Kit(T3001). N different PCN-plasmid were tested with the same main inverter plasmid. The expression of sfGFP from dual plasmid system with different PCN was individually measured at a gradient of 20 aTc concentrations on the BioTek (Synergy H1) plate reader in H medium. Three biological replicates were used for each system, i.e., three different single colony were picked up and grown in 2 mL H media without aTc for approximately 18 hours. Then, 100 μL of overnight cell culture was used and diluted into 2 mL fresh H media before transferring to a 96 well plate where each of the middle of 10 x 6 wells contains 1 μL of diluted cell culture and 200 μL fresh H media with corresponding aTc concentration.

### Growth Rate Measurements

Growth rates of individual constructs were measured on the BioTek (Synergy H1) plate reader. Overnight cultures of cells grown to saturation were serially diluted by a factor of 100,000 into well-mixed 200 μL aliquots of media containing the appropriate antibiotics and inducer concentrations. Cells were grown at 37°C for 36 h while to OD_600_ and fluorescence (if appropriate) was recorded every 3 min. Growth rates were calculated from the resultant time series data by averaging a window of ± 6 min around the maximal growth rate, calculated as a discrete difference of OD_600_ readings between time points.

### Gene Expression Measurements

The expression of green fluorescent protein (sfGFP) was used a measure of gene expression in this work. Fluorescence measurements were taken on cells prepared as described above as they grew. Gene expression was calculated as follows - the average fluorescence in a small window surrounding the maximal growth rate was calculated and divided by the integral of the OD_600_ in the same window. This can be interpreted as a the fluorescence (ie gene expression) per cell and should be independent of the growth rate since we divide by the integral of the OD_600_. Six replicates of each inducer concentration were taken and the average and standard deviation of this measure of gene expression were reported.

### Violacein Production Measurements

Cultures of single colonies hosting plasmids containing the violacein biosynthesis pathway were grown overnight in Terrific Broth with appropriate antibiotic. First, 200 μL of liquid culture was centrifuged and cells were resuspended in 50 μL of water. Then, 450 μL of methanol was added to the resuspended cells and they were shaken for 1 h at room temperature. Cells were spun down and 200 μL of the supernatant was transferred to a 96-well plate where the absorbance at 650 nm was used to quantify violacein production.

### Single cell microscopy

The single cell data was collected within a month over 8 subsequent experiments with varying aTc concentrations. Each experiment was run for 1300 to 5000 min, corresponding to at least 20 to 80 generations. All experiments were started from the same stock bacterial colony, initially grown to OD_600_ 0.5 at 10ng mL^−1^ aTc and kept refrigerated. In order to start the experiment, 3 μL of the stock colony was disperse in 3mL of H media with plasmid selecting antibiotic carbenicillin and aTc. The cells were grown to OD_600_ 0.5 ± 0.1, concentrated by centrifugation and inserted in plasma cleaned microfluidic chips.

After entering the chip bacteria populated and grew in 60 μm long, 7.5 μm wide and 1.05 μm thick chambers connected to a supply channel flowing fresh media pressurized to 4 psi [22]. On top of H media, carbenicillin and aTc, 0.1 gL^−1^ of bovine serum albumin was added to the media to improve evacuation of cells from the supply channel [42].

Single cell were resolved a by epi-fluorescence microscopy of sf-GFP with a 100 x, 1.4NA apochromat Leica objective. Single cell detection and tracking enabled measurement of cell dimensions, intensity and growth rate overtime. Cells can be classified in three categories: (1) non dividing dark cells (loss of antibiotic resistance due to plasmid loss), (2) healthy cells and (3) non growing bright cells (likely due to plasmid overload). Whereas in bulk pathological cells of type (1) or (3) would quickly be overgrown by healthy cells, in chambers, we find that they can get stuck at the back or even populate the whole chamber, rendering the chamber unusable when the cells eventually die. This is particularly true for non dividing dark cells at low aTc concentration. By using 7.5 μm wide chambers, we allow many cells to grow side by side, which increases the chances of healthy cells to repopulate the chambers. However, this is not enough to maintain healthy chambers for more than a few hours at low aTc concentration. For aTc concentrations of 1 ng mL^−1^ and above a majority of chambers last for the whole experiment time of 5000 min. We base intensity measurements on growing cells only (growth rate larger than 1/3 of mean growth rate) to prevent over representation of non growing cells that can reach up to 40% of the detected cells (Fig. 3F). Selected cells are highlighted in supplementary movie.

Due to the rapid loss of chambers we are unable to measured a stabilized fluorescence level for aTc concentration below 1 ng mL^−1^. However, we can still observe that the aTc concentration promote plasmid replication. Indeed, the chamber survive longer at higher aTc concentration (plasmid loss is less frequent) and the average fluorescence increases with aTc. For aTc concentration of 1 ng mL^−1^ to 100ng mL^−1^ time dependent fluorescence and growth rate curves (Figure S2) stabilize after about 600 min (i.e. 12 generations) and are expected to accurately represent PCN in exponential growth condition.

All experiments and cell cultures were performed at 37°C.

## ACKNOWLEDGMENTS

We wish to thank David Specht for helpful discussions and critical review of the manuscript. We also thank members of the Lambert laboratory for providing feed-back on the manuscript. This work was supported by NIH funding under 1R35 GM133759 Maximizing Investigators’ Research Award (MIRA).

## Supplementary Materials

### MOTHER MACHINE

#### Data selection

In order to discard pathological cells as well as detection errors the following threshold are used to select data.

1. 0.7 μm< *D* <4.3 μm
2. 0.005 min^−1^< growth rate <0.04 min^−1^
3. 1.7 μm< *L* <17 μm
4. intensity> 122 A.U.

The intensity threshold is set just a few unit above the average background (110-120). The most restrictive conditions is the one on growth rate. It is used to set apart non growing cells as well as cells that can not be tracked overtime due to detection errors. Supplementary figure S1 shows histogram of cell properties before and after selection for one of the worst cases at 0.01 ng mL^−1^. The peak at 0 correspond to cells without measured growth rate. Detection and tracking error may lead to cell tracks which length decreases overtime leading to negative growth rate.

#### Cell morphology

In supplementary figure S2, we compare histograms of cells morphology for all aTc conditions. We observe that the cells are longer at 10 ng mL^−1^ aTc than 0.01 ng mL^−1^. However, the volume - surface area ratio is conserved.

#### Stabilization of PCN

In S3 we plot cell properties as a function of time for each aTc concentration.

The number of detected cells quickly increases when cells populate the chambers. Then, the total number of detected cells decreases for all aTc conditions. This is largely due to chambers being filled by pathological cells that upon death block their chamber. Pathological cells mostly suffer from plasmid loss at low aTc while plasmid overload dominates at high aTc. We observe that plasmid loss is more prone to block chambers. Indeed, the more aTc, i.e. the more PCN, the longer lived are the chambers.

For aTc concentration of 1ng mL^−1^ and above, we observe a stabilization of the average cell intensity after about 600 minutes (about 12 generations). We also observe a stabilization in growth rate around this time. Only the data points past this 600 min threshold are used in Fig. 3B and 3C of the main text, as well as supplementary figure 4, for aTc conditions of 1 ng mL^−1^ and above.

For aTc concentration of 0.1 ng mL^−1^ and below, plasmid loss is too frequent, we are not able to measure a stabilized average cell intensity. We use all time points available for those concentrations in other plots across the manuscript.

#### Growth rate

Growth rate measurement particularly lack of precision for the low aTc conditions. Considering aTc concentration of 1 ng mL^−1^ and above we observe a constant growth rate with a 10% decrease for the highest aTc condition of 100 ng mL^−1^.

#### PCN distribution

In supplementary figure S5, we plot fold change histograms and corresponding gamma distributions:

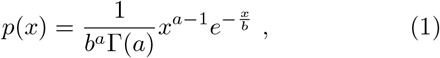

with Γ the gamma function, 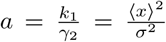 the mean number of burst per cycle (Friedman PRL 2006) and 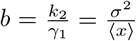 the mean burst size.

### MASSIVELY PARALLELIZED ASSAY

#### No Relationship Between Priming Promoter Strength and Plasmid Copy Number

Given that we saw a clear relation between promoter de-repression and plasmid copy number (Fig. 1D) and that high-copy number plasmids had similar origins of replication, we were surprised that we could not find a clear relation between the predicted strength [14] of the promoter controlling RNA-p and the plasmid copy number (Fig. S6).

### TABLE I

Table S1: Promoter Sequences, Next-generation sequencing counts, Relative Growth Rates, Predicted Plasmid Copy Numbers, and Predicted Promoter Strength for priming RNA variants used in this work

### TABLE II

Table S2: Promoter Sequences, Next-generation sequencing counts, Relative Growth Rates, Predicted Plasmid Copy Numbers, and Predicted Promoter Strength for inhibitory RNA variants used in this work

### TABLE III

Table S3: Sequencing Counts at each time point for priming RNA variants

**Fig. S1.**
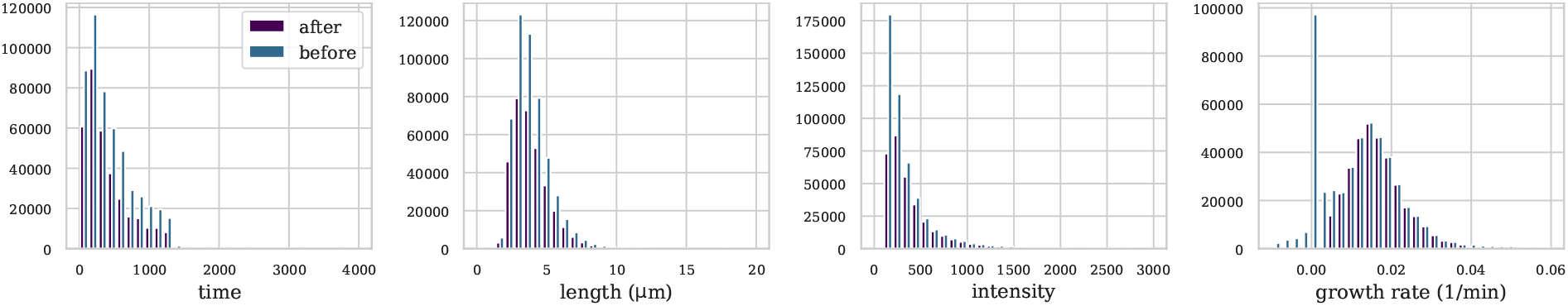
Count of cells as a function of time, length, fluorescence intensity, and growth rate before and after selection for one of the most difficult condition at 0.01 ng mL^−1^ aTc.

**Fig. S2.**
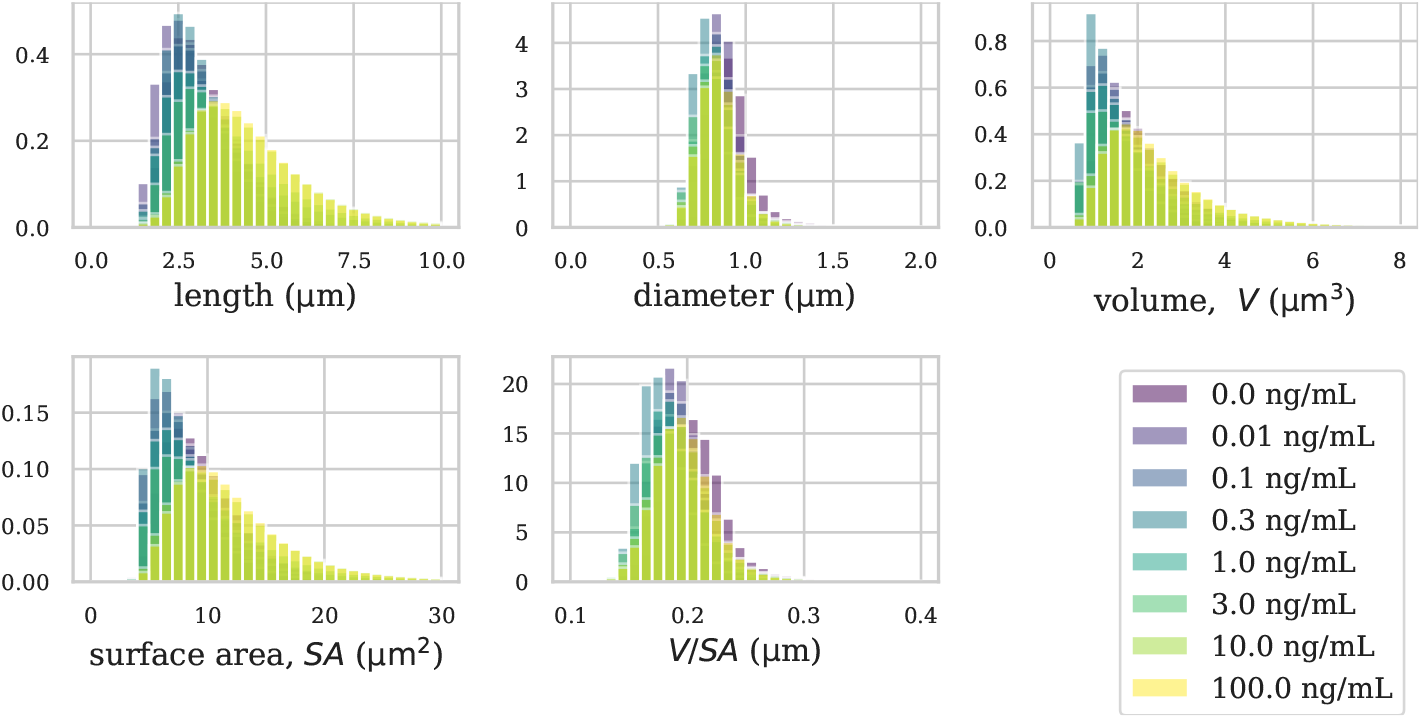
Increase of cell length with aTc concentration, i.e. plasmid copy number, but conserved volume/surface area ratio.

**Fig. S3.**
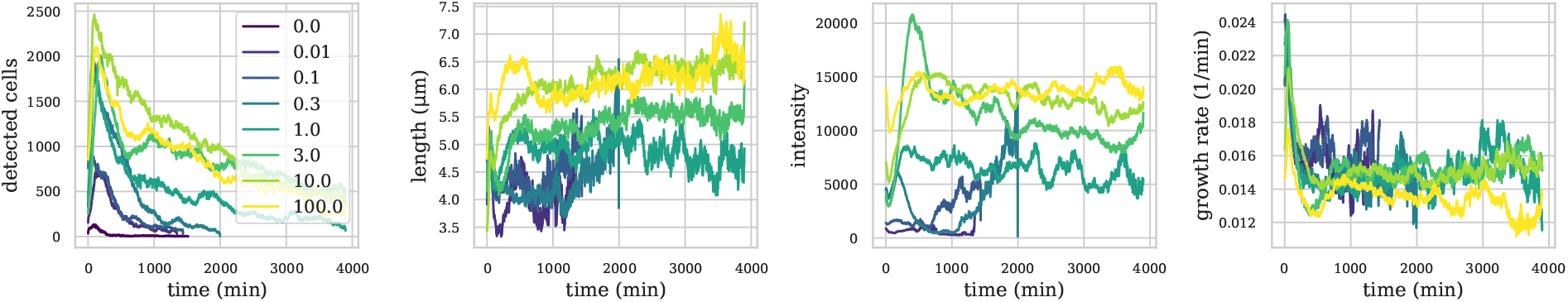
Time dependent measurement of the number of detected cells, their averaged length, fluorescence and growth rate.

**Fig. S4.**
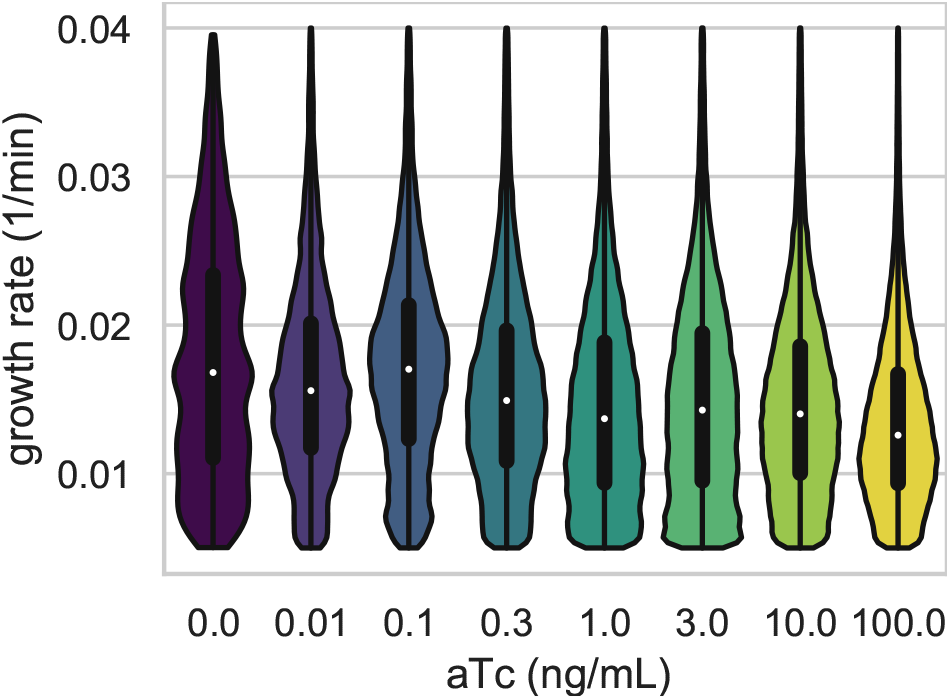
Violin plots illustrating the distribution of growth rates for all aTc concentrations after selection (c.f. supplementary figure 1).

**Fig. S5.**
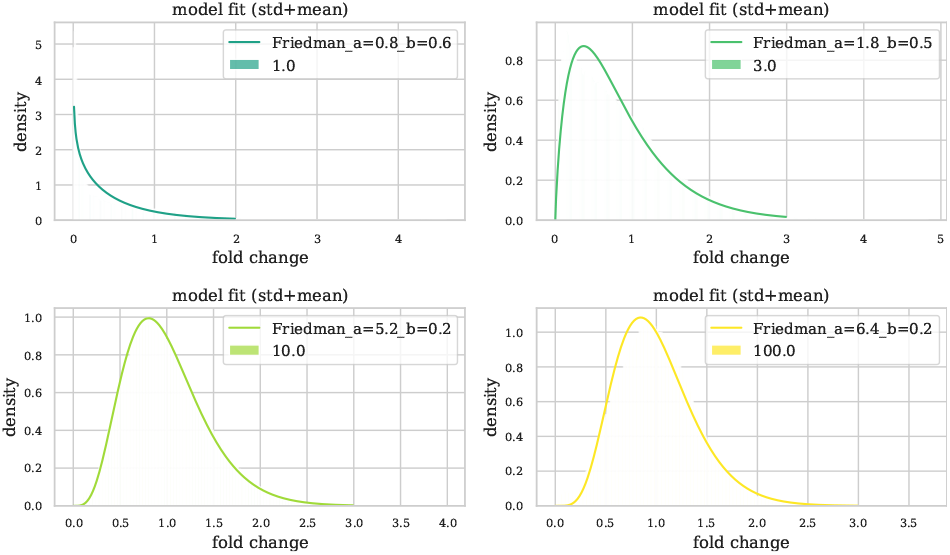
Fit of fold change histograms for stabilized movies at 1, 3, 10 and 100ng mL^−1^ aTc.

**Fig. S6.**
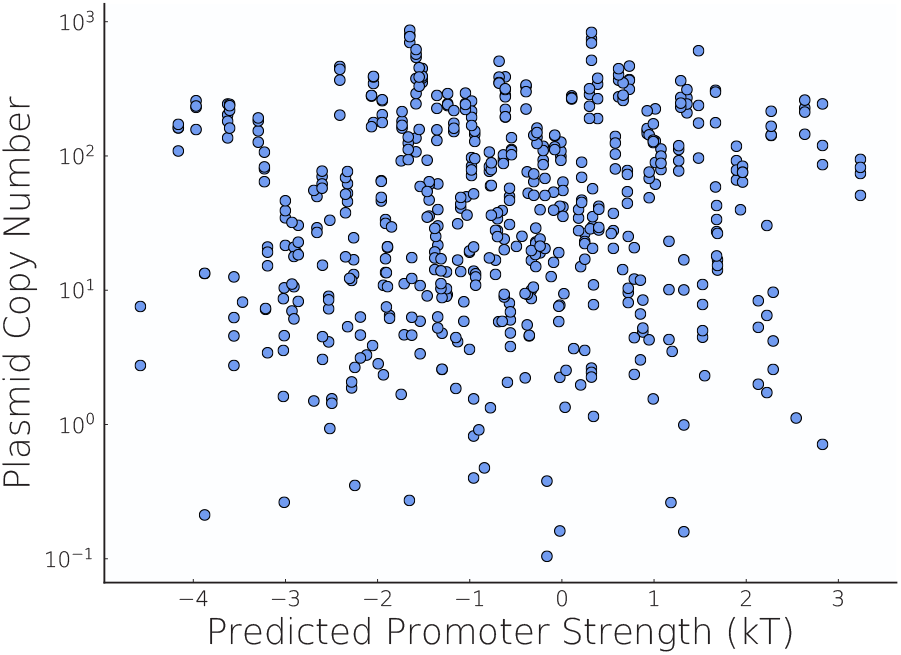
Priming Promoter Strength vs Plasmid Copy Number

## References

[1] A. P. Togna, M. L. Shuler, and D. B. Wilson, Effects of Plasmid Copy Number and Run-away Plasmid Replication on Overproduction and Excretion of -Lactamase from Escherichia coli, Biotechnology Progress 9, 31 (1993), _eprint: https://aiche.onlinelibrary.wiley.com/doi/pdf/10.1021/bp00019a005d

[2] K. Nordström and B. E. Uhlin, Runaway-replication plasmids as tools to produce large quantities of proteins from cloned genes in bacteria, Bio/Technology (Nature Publishing Company) 10, 661 (1992).

[3] M. Dwidar and Y. Yokobayashi, Riboswitch Signal Amplification by Controlling Plasmid Copy Number, ACS Synthetic Biology 8, 245 (2019), publisher: American Chemical Society.

[4] R. U. Sheth, S. S. Yim, F. L. Wu, and H. H. Wang, Multiplex recording of cellular events over time on CRISPR biological tape, Science 358, 1457 (2017), publisher: American Association for the Advancement of Science Section: Report.

[5] L. Potvin-Trottier, N. D. Lord, G. Vinnicombe, and J. Paulsson, Synchronous long-term oscillations in a synthetic gene circuit, Nature 538, 514 (2016).

[6] B. Shao, J. Rammohan, D. A. Anderson, N. Alperovich, D. Ross, and C. A. Voigt, Single-cell measurement of plasmid copy number and promoter activity, Nature Communications 12, 1475 (2021).

[7] J. C. Diaz Ricci and M. E. Hernández, Plasmid Effects on *Escherichia coli* Metabolism, Critical Reviews in Biotechnology 20, 79 (2000).

[8] A. S. Karim, K. A. Curran, and H. S. Alper, Characterization of plasmid burden and copy number in Saccharomyces cerevisiae for optimization of metabolic engineering applications, FEMS Yeast Research 13, 107 (2013), _eprint: https://onlinelibrary.wiley.com/doi/pdf/10.1111/1567-1364.12016.

[9] D. S.-W. Ow, P. M. Nissom, R. Philp, S. K.-W. Oh, and M. G.-S. Yap, Global transcriptional analysis of metabolic burden due to plasmid maintenance in Escherichia coli DH5 during batch fermentation, Enzyme and Microbial Technology 39, 391 (2006).

[10] W. E. Bentley, N. Mirjalili, D. C. Andersen, R. H. Davis, and D. S. Kompala, Plasmid-encoded protein: The principal factor in the “metabolic burden” associated with recombinant bacteria, Biotechnology and Bioengineering 35, 668 (1990), _eprint: https://onlinelibrary.wiley.com/doi/pdf/10.1002/bit.260350704

[11] M. Pasini, A. Fernández-Castané, A. Jaramillo, C. de Mas, G. Caminal, and P. Ferrer, Using promoter libraries to reduce metabolic burden due to plasmid-encoded proteins in recombinant Escherichia coli, New Biotechnology 33, 78 (2016).

[12] A. Rozkov, C. A. Avignone-Rossa, P. F. Ertl, P. Jones, R. D. O’Kennedy, J. J. Smith, J. W. Dale, and M. E. Bushell, Characterization of the metabolic burden on Escherichia coli DH1 cells imposed by the presence of a plasmid containing a gene therapy sequence, Biotechnology and Bioengineering 88, 909 (2004).

[13] F. M. Weinert, R. C. Brewster, M. Rydenfelt, R. Phillips, and W. K. Kegel, Scaling of Gene Expression with Transcription-Factor Fugacity, Physical review letters 113, 258101 (2014).

[14] R. C. Brewster, F. M. Weinert, H. G. Garcia, D. Song, M. Rydenfelt, and R. Phillips, The Transcription Factor Titration Effect Dictates Level of Gene Expression, Cell 156, 1312 (2014).

[15] J. Paulsson, K. Nordström, and M. Ehrenberg, Requirements for Rapid Plasmid ColE1 Copy Number Adjustments: A Mathematical Model of Inhibition Modes and RNA Turnover Rates, Plasmid 39, 215 (1998).

[16] G. del Solar, R. Giraldo, M. J. Ruiz-Echevarría, M. Espinosa, and R. Díaz-Orejas, Replication and Control of Circular Bacterial Plasmids, Microbiology and Molecular Biology Reviews 62, 434 (1998).

[17] J.-i. Tomizawa, Control of cole1 plasmid replication: Initial interaction of RNA I and the primer transcript is reversible, Cell 40, 527 (1985).

[18] J. Paulsson and M. Ehrenberg, Noise in a minimal regulatory network: plasmid copy number control, Quarterly Reviews of Biophysics 34, 1 (2001), publisher: Cambridge University Press.

[19] J. Paulsson, Multileveled selection on plasmid replication., Genetics 161, 1373 (2002).

[20] M. Plotka, M. Wozniak, and T. Kaczorowski, Quantification of Plasmid Copy Number with Single Colour Droplet Digital PCR, PLoS ONE 12, 10.1371/journal.pone.0169846 (2017).

[21] P. Wang, L. Robert, J. Pelletier, W. L. Dang, F. Taddei, A. Wright, and S. Jun, Robust Growth of Escherichia coli, Current Biology 20, 1099 (2010).

[22] G. Lambert and E. Kussell, Memory and Fitness Optimization of Bacteria under Fluctuating Environments, PLoS Genet 10, e1004556 (2014).

[23] D. A. Specht, Y. Xu, and G. Lambert, Massively parallel CRISPRi assays reveal concealed thermodynamic determinants of dCas12a binding, Proceedings of the National Academy of Sciences 117, 11274 (2020).

[24] H. C. Lichstein and V. F. Van De Sand, Violacein, an Antibiotic Pigment Produced by Chromobacterium Violaceum, The Journal of Infectious Diseases 76, 47 (1945), publisher: Oxford Academic.

[25] M. Kholany, P. Tréebulle, M. Martins, S. P. Ventura, J. Nicaud, and J. A. Coutinho, Extraction and purification of violacein from *Yarrowia lipolytica* cells using aqueous solutions of surfactants, Journal of Chemical Technology & Biotechnology, jctb.6297 (2019).

[26] D. G. Gibson, L. Young, R.-Y. Chuang, J. C. Venter, C. A. Hutchison, and H. O. Smith, Enzymatic assembly of DNA molecules up to several hundred kilobases, Nature Methods 6, 343 (2009).

[27] M. R. Atkinson, M. A. Savageau, J. T. Myers, and A. J. Ninfa, Development of Genetic Circuitry Exhibiting Toggle Switch or Oscillatory Behavior in Escherichia coli, Cell 113, 597 (2003).

[28] Y. Mileyko, R. I. Joh, and J. S. Weitz, Small-scale copy number variation and large-scale changes in gene expression, Proceedings of the National Academy of Sciences 105, 16659 (2008).

[29] T. Wang, N. Tague, S. A. Whelan, and M. J. Dunlop, Programmable gene regulation for metabolic engineering using decoy transcription factor binding sites, Nucleic Acids Research 49, 1163 (2021).

[30] B. Wang, F. Guo, S.-H. Dong, and H. Zhao, Activation of silent biosynthetic gene clusters using transcription factor decoys, Nature Chemical Biology 15, 111 (2019).

[31] M. Z. Ali and R. C. Brewster, Controlling gene expression timing through gene regulatory architecture, bioRxiv, 2021.04.09.439163 (2021).

[32] X. Wan, F. Pinto, L. Yu, and B. Wang, Synthetic protein-binding DNA sponge as a tool to tune gene expression and mitigate protein toxicity, Nature Communications 11, 5961 (2020).

[33] S. Zhang and C. A. Voigt, Engineered dCas9 with reduced toxicity in bacteria: implications for genetic circuit design, Nucleic Acids Research 46, 11115 (2018).

[34] C. Tan, P. Marguet, and L. You, Emergent bistability by a growth-modulating positive feedback circuit, Nature Chemical Biology 5, 842 (2009).

[35] Y. Qian, H.-H. Huang, J. I. Jiménez, and D. Del Vecchio, Resource Competition Shapes the Response of Genetic Circuits, ACS Synthetic Biology 6, 1263 (2017).

[36] F. Ceroni, R. Algar, G.-B. Stan, and T. Ellis, Quantifying cellular capacity identifies gene expression designs with reduced burden, Nature Methods 12, 415 (2015).

[37] J. Tomizawa and T. Itoh, Plasmid ColE1 incompatibility determined by interaction of RNA I with primer transcript., Proceedings of the National Academy of Sciences 78, 6096 (1981).

[38] J. Rodríguez-Beltrán, J. DelaFuente, R. León-Sampedro, R. C. MacLean, and San Millán, Beyond horizontal gene transfer: the role of plasmids in bacterial evolution, Nature Reviews Microbiology 19, 347 (2021).

[39] J. Ilhan, A. Kupczok, C. Woehle, T. Wein, N. F. Hülter, P. Rosenstiel, G. Landan, I. Mizrahi, and T. Dagan, Segregational Drift and the Interplay between Plasmid Copy Number and Evolvability, Molecular Biology and Evolution 36, 472 (2019), publisher: Oxford Academic.

[40] T. Vogwill and R. C. MacLean, The genetic basis of the fitness costs of antimicrobial resistance: a meta-analysis approach, Evolutionary Applications 8, 284 (2015), _eprint: https://onlinelibrary.wiley.com/doi/pdf/10.1111/eva.12202.

[41] T. Wein, N. F. Hülter, I. Mizrahi, and T. Dagan, Emergence of plasmid stability under non-selective conditions maintains antibiotic resistance, Nature Communications 10, 2595 (2019).

[42] D. Yang, A. D. Jennings, E. Borrego, S. T. Retterer, and J. Männik, Analysis of factors limiting bacterial growth in pdms mother machine devices, Frontiers in Microbiology 9, 871 (2018).

